# Input–Output Slope Curve Estimation in Neural Stimulation Based on Optimal Sampling Principles

**DOI:** 10.1101/2021.02.16.431436

**Authors:** Seyed Mohammad Mahdi Alavi, Stefan M. Goetz, Mehrdad Saif

## Abstract

This paper discusses some of the practical limitations and issues, which exist for the input–output (IO) slope curve estimation (SCE) in neural, brain and spinal, stimulation techniques. The drawbacks of the SCE techniques by using existing uniform sampling and Fisher-information-based optimal IO curve estimation (FO-IOCE) methods are elaborated. A novel IO SCE technique is proposed with a modified sampling strategy and stopping rule which improve the SCE performance compared to these methods. The effectiveness of the proposed IO SCE is tested on 1000 simulation runs in transcranial magnetic stimulation (TMS), with a realistic model of motor evoked potentials (MEPs). The results show that the proposed IO SCE method successfully satisfies the stopping rule, before reaching the maximum number of TMS pulses in 79.5% of runs, while the estimation based on the uniform sampling technique never converges and satisfies the stopping rule. At the time of successful termination, the proposed IO SCE method decreases the 95th percentile (mean value in the parentheses) of the absolute relative estimation errors (AREs) of the slope curve parameters up to 7.45% (2.2%), with only 18 additional pulses on average compared to that of the FO-IOCE technique. It also decreases the 95th percentile (mean value in the parentheses) of the AREs of the IO slope curve parameters up to 59.33% (16.71%), compared to that of the uniform sampling method. The proposed IO SCE also identifies the peak slope with higher accuracy, with the 95th percentile (mean value in the parentheses) of AREs reduced by up to 9.96% (2.01%) compared to that of the FO-IOCE method, and by up to 46.29% (13.13%) compared to that of the uniform sampling method.

## I. Introduction

**V**Arious stimulation techniques, such as transcranial electrical stimulation (TES) and transcranial magnetic stimulation (TMS) allow relatively selective activation of neurons in the central nervous system [1], [2]. If administered to the motor system, a response in the form of a motor evoked potential (MEP) can be detected through electromyography (EMG) in a peripheral muscle that corresponds to the activated representation. Neural motor recruitment or input–output (IO) curves represent the relationship between stimulus strength and MEP size in response to such neural stimulation with TES or TMS in the motor cortex or the spine [3]–[5].

IO curves reflect the neural excitability of a stimulation target, which is affected by neuromodulators, incoming endogenous signals, the brain state, as well as a variety of disorders. Extensive research is covering the relationship between the IO curve parameters and medical conditions. For instance, IO curves can be used as biomarkers for movement disorders [6], [7], epilepsy [8], dementia [9], stroke [10], [11], idiopathic normal pressure hydrocephalus (iNPH) [12], neurodegenerative diseases [13], and various other neurological conditions [14]–[18].

The IO curve reflects the entire stimulus–response behavior from weakest stimuli with hardly any response to strongest pulses where the response saturates, including aspects such as maximum recruitment and physiological variability from different sites and mechanisms. For very weak stimuli, the IO curve forms a flat plateau as the responses vanish in the technical noise floor, which can be reduced by better detection techniques [19]. For very strong stimuli, the curve flattens on the upper side as the responses saturate. In between, an S-shaped region describes the transition with a well-defined slope as the tangent or derivative of the IO curve.

Particularly the slope of the IO curve, i.e., this average growth of recruitment with increasing stimulation strength, is in the focus of most IO-curve biomarker applications. The slope of the IO curve represents a general measure of the cortical excitability [20]. It gives an indication of the distribution of excitability in the corticospinal projection and reflects the level of synaptic connectivity and plasticity in the cerebral cortex and the spinal cord [21]–[25]. The narrower the excitability distribution, the larger the available connections, which tends to result in a steeper slope of the IO curve, and vice versa. For instance, in [12], the IO curve of a number of iNPH patients were acquired by TMS before and after cerebrospinal fluid drainage through lumbar puncture. The results show significant increase of the slope of the IO curve after drainage in iNPH patients who had improved gait performance. By further neurophysiological studies, it was concluded that the observed steeper slope in the iNPH patients with improved gait performance is possibly due to a higher cortical recovery potential by better preserved synaptic connectivity and plasticity.

The slope is not constant in the S-shaped region of the IO curve and changes with respect to the pulse amplitude. The change of the slope is understandable in the light that along the IO curve different neuron populations join the pool of activated neurons, presumably starting with various types of interneurons and expanding to pyramidal neurons for stronger pulses [26]. The slope curve determines the pointwise slope of the IO curve at any given pulse amplitude, which can be modelled by a normal function in general. Studies confirm that the slope curve depends on age, sex, or other neurophysiological conditions. For instance, the results in [17] show the shift of the slope curve to the right for older subjects. The peak of the slope curve’s normal function reflects the gain of maximum corticospinal output from the TMS pulse, [21], and has been identified as a neurophysiological biomarker. The peak slope of the IO curve increases with increasing the tonic background activity up to 4 – 7 times the value at rest for the first dorsal interosseous and tibialis anterior muscles [22]. Significant differences were also reported between the peak slope of the affected and unaffected hemispheres in stroke patients [11].

Traditionally, TMS IO curves are estimated through uniform sampling [10], [11], [27]–[33]. A number of uniformly distributed TMS pulses are administered to the subject. Whereas early studies even used an order, e.g., from weak to strong stimuli, strong correlation between subsequent responses, which shows up as hysteresis, strongly advises against that [34], [35]. Instead, a random order is recommended. The MEP value is measured after each pulse, the data set of the stimulus–response pairs constructed, and the IO curve estimated by using a curve fitting algorithm. In order to address the variabilities, TMS pulses are sometimes repeated multiple times for each stimulation strength, and the average of MEPs is taken for curve fitting.

A recently developed method suggested sequential estimation of the TMS IO curve and its parameters using Fisher information matrix (FIM) optimization to not only speed up the procedure, but also increase accuracy and independence from arbitrary sampling decisions [36]. The FIM-based optimal IO curve estimation (FO-IOCE) method also administers the next stimulus such that it achieves faster convergence of the IO curve estimation with a desired level of accuracy, compared to simpler uniform or arbitrary sampling. In consequence, such advanced FO-IOCE method can reduce estimation errors up to two-fold and requires ten-times fewer stimuli than uniform sampling [36].

In both uniform sampling and FO-IOCE methods, the overall slope of the IO curve is estimated within the estimation process of the IO curve. The slope curve is subsequently estimated by taking the derivative of the estimated IO curve.

However, the conventional uniform sampling and FO-IOCE methods suffer from a number of drawbacks. Despite simplicity, the main drawback of the uniform sampling method is that it does not provide any information about the number of TMS pulses, their repetition, and strengths necessary to achieve a satisfactory level of estimation accuracy as quickly as possible, nor about the current expected prediction error. These parameters are usually chosen empirically in the uniform sampling literature and often disregard individual variability [10], [11], [27]–[33]. The uniform sampling method is also not optimal in the sense that many TMS pulses might be administered that do not necessarily improve the estimation accuracy or precision.

The IO curve is sigmoidal, and the slope curve is a normal function. The locations of maximum information are different for these two functions although there are some overlaps. Thus, the FO-IOCE method is not optimal for slope curve estimation (SCE).

This paper elaborates on the practical limitations and challenging issues associated with the IO SCE problem. Subsequently, a novel sub-optimal method is developed for the slope curve estimation (SCE) in neural stimulation. The proposed method builds upon previous Fisher-information-based estimation but introduces a novel sampling strategy and stopping rule, which improve the estimation performance compared to the FO-IOCE method. The effectiveness of the proposed IO SCE is tested on 1000 simulation runs in transcranial magnetic stimulation (TMS), and the results are discussed and compared with the FO-IOCE and uniform sampling methods.

Based on these statements, the main contributions of this paper are two:

- Explaining optimal sampling for IO curve and slope curve estimation, and
- Developing a novel IO SCE method for neural stimulation techniques, with modified sampling strategy and stopping rule, which improve the estimation performance compared to the FO-IOCE and uniform sampling methods.

The rest of the paper is outlined as follows. In Section II, the IO curve and its derivative models are reviewed. The problem of this paper is clearly formulated. Some practical limitations and issues in the estimation of the slope curve are also discussed. In Section III, the proposed IO SCE method is elaborated. In Section IV, the effectiveness of the proposed IO SCE method is evaluated and compared with the uniform sampling and FO-IOCE methods.

## II. Some Basic Information

### A. Models

Figure 1 shows a typical data set of stimulus–response pairs in TMS. The IO curve, *y*(*x*), is obtained by fitting a sigmoid model to this data set. The majority of the literature [10], [11], [27]–[31], [36], employ the four-parameter identifiable [37] sigmoid model

**Fig. 1:**
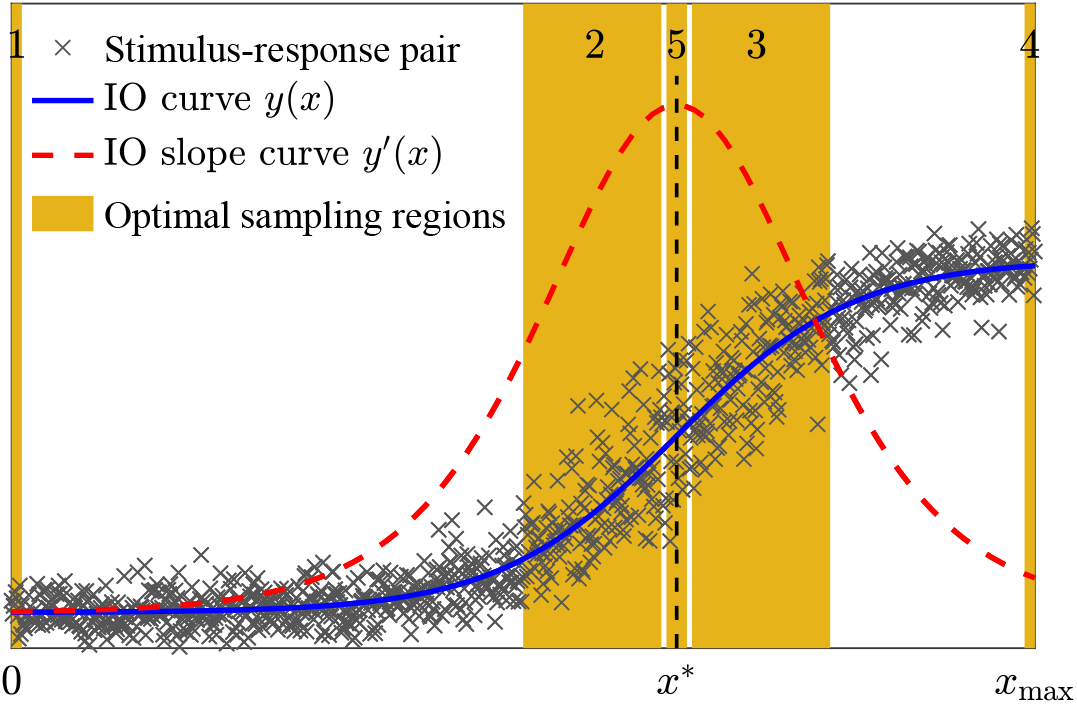
A typical data set of stimulus–response pairs in neural stimulation applications, the IO curve *y*, and the IO slope curve *y ′*, and optimal sampling regions. The slope curve *y ′* (*x*) was scaled for a better visualization. Based on optimal sampling principles, sampling from the Areas 1, 2, 3, and 4 is required for the estimation of the IO curve *y*(*x*). For the estimation of the normal function *y ′* (*x*), sampling from the peak-point Region 5, and one of either Regions 2 or 3 is enough.

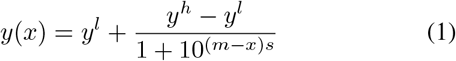

where *x* is the TMS stimulus, *y* denotes a characteristic of MEP such as its peak-to-peak value, area under the curve (AUC), root mean square or latency [33], and *y*^*l*^, *y*^*h*^, *m*, and *s* are the low-side plateau, high-side plateau, midpoint, and slope parameters, respectively. The peak-to-peak MEP at *x* = *m* is the mean of the low and up plateaus, i.e.,

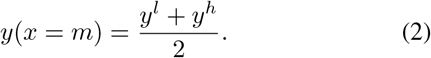

The parameter vector ***θ*** is defined as

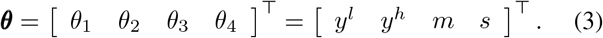

Figure 1 also shows the IO slope curve *y′ (x)*, which is the derivative of *y* with respect to the neural stimulus *x*, given by

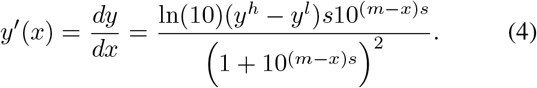

The IO slope curve *y′* (*x*) denotes the slope of the IO curve at the stimulus strength *x*. As it is seen, the slope of the IO curve varies with respect to *x*, as a normal function, with a maximum (peak) slope at *x*\* = *m*, where the second derivative reaches zero. Consequently, the peak slope is given by

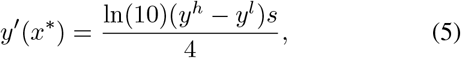

confirming that the peak slope is proportional to the slope parameter *s*, with the gain of ln(10)(*y*^*h*^ *− y*^*l*^)*/*4 for the IO curve model (1).

Note that *s* is the slope parameter, and *y′* is the slope curve. The value of *s* determines the overall slope of the IO curve; the larger the *s* value, the steeper the slope of the IO curve. In contrast, *y′* (*x*) provides the pointwise slope information, i.e, the slope of the IO curve at the stimulus strength *x*.

### B. Problem statement

The main problem in this paper is to reliably estimate the IO slope curve *y*′ and the peak slope (*x*\*, *y′* (*x*\*)), with a satisfactory level of accuracy and as few as possible TMS pulses.

### C. Practical limitations & challenging issues

In addition to the limitation on the test duration, which demands for estimation methods with as few as possible TMS pulses, there is another main practical limitation that there is no tool for the direct measurement of *y′* at a pulse strength *x*. More importantly, due to the large noise and variabilities which exist along both x and y axes [38], the IO curve derivative at a certain pulse strength *x* cannot be measured by applying two TMS pulses, one at *x* and the other one at *x* + Δ*x*, and using the numerical differentiation formula [39], [40]

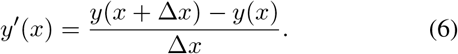

The only measurable data set is the stimulus–response pairs of the IO curve, including the baseline data, in the form of

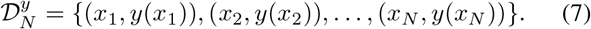

## III. The Proposed Slope Curve Estimator

Since 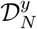 is the only measurable data set, a possible IO SCE method is to firstly estimate the parameter vector ***θ*** through the IO curve estimation, and then calculate the slope curve *y*′ by using the most recent estimate of the parameter vector and (4).

The FO-IOCE method estimates the parameter vector ***θ***, by using the baseline data from Region 1, as well as the TMS stimuli, which are optimally chosen from Regions 2, 3, and 4, as shown in Figure 1. The baseline data is the EMG data in the absence of TMS pulse for *x* = 0.

The slope curve is like a normal function. It is shown in [41] that sampling from the peak-point region 5, and one of either regions 2 or 3 is required and enough for the optimal estimation of the normal function.

Based on these statements, the FO-IOCE method [36] is not optimal for the IO SCE, since it does not take samples from the peak-point Region 5.

In this section, an IO SCE technique is presented, which is a modified version of the FO-IOCE idea, guaranteeing sampling from the peak-point Region 5, and one of either Regions 2 or 3, improving the estimation performance.

Figure 2 describes the proposed IO SCE, which starts with the measurement of the *n*_*base*_ baseline data, applying three randomly TMS pulses, measuring their MEPs, and constructing the initial data set 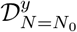. An initial estimate of the parameter vector, 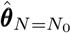 is obtained by fitting the four-parameter sigmoid model (1) to the data set of 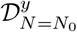. The next TMS pulse, *x*_*N*+1_ is computed by solving the following FIM optimization problem,

**Fig. 2:**
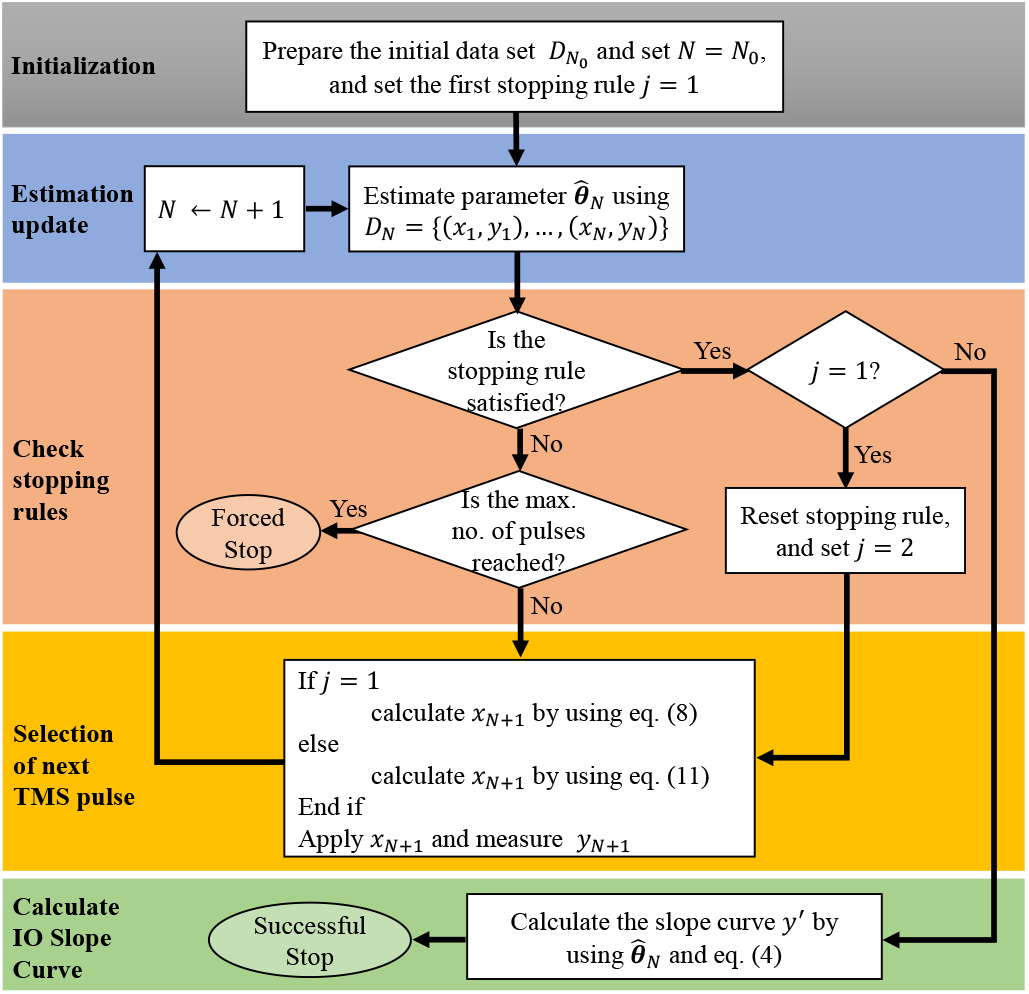
Flowchart of the proposed input-output (IO) slope curve estimation (SCE) method based on optimal sampling principles.

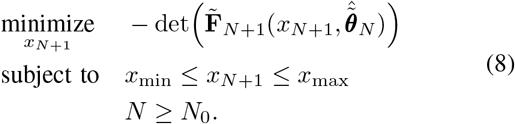

where 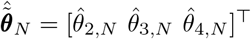, and

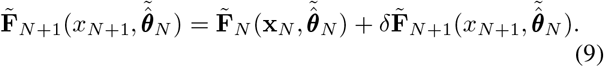

The matrices 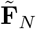 and 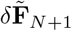 are given in Appendix A. The solution of (8) helps optimal estimation of the parameter vector by stimulation of TMS pulses from the locations of maximum information, i.e., Regions 2, 3, and 4. The next TMS pulse *x*_*N*+1_ is stimulated, and the corresponding MEP, *y*_*N*+1_ is measured. The data set of 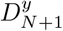 is obtained. The curve fitting is re-run, and 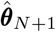 is achieved. This process continues until the convergence criterion

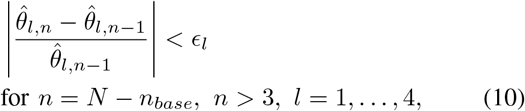

is satisfied for *T* successive times. Satisfaction of (10) for *T* successive times refers to as the stopping rule. The parameter *ϵ*_*l*_ is the convergence tolerance. The estimation precision is adjustable by using *T* and *ϵ*_*l*_ values. The larger *T* and the smaller *ϵ* _*l*_ result in more accurate estimation, at the cost of more number of TMS stimuli. Therefore, there are trade-offs. Assume that the the stopping rule is satisfied at 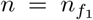. Up to this point, the IO curve is estimated by applying TMS stimuli from the regions 1 to 4. Hereafter, the proposed SCE method continues estimation by applying more TMS pulses, which target stimulation at the peak-point Region 5 per

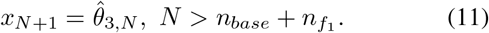

The stopping rule is reset at 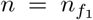 and re-evaluated from scratch. Once it is satisfied for the second time at 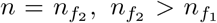, the estimation successfully terminates, and the IO slope curve is estimated by using (4) and 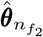. If the stopping rule is not satisfied in any of two cases, for the first or second time, TMS continues until a forced stop, when the maximum number of pulses is administered.

The source code of the proposed IO SCE method is available from [42].

## IV. Results

The effectiveness of the proposed method is tested synthetically in Matlab over 1000 simulation runs during TMS. In this study, the MEP’s peak-to-peak data is used representing the *y* parameter in (1). This simulation employs a detailed statistical model derived from experiments [43], which reproduces the intricate statistical distributions observed in measurements [44], [45] by generating near-realistic stimulus–response pairs in non-invasive brain stimulation techniques, considering neurophysiological and electromagnetic uncertainties and variabilities such as trial-to-trial variabilities, excitability fluctuations, variability in the neural and muscular pathways, electric field’s amplitude and focality, and physiological and measurement noise. If the variations and uncertainties are large, it takes longer for the estimation process to satisfy the stopping rule and stop. Keeping the coil position and orientation, and subject’s head fixed during the test helps reduce the variations and uncertainties and expedite the estimation process.

In each run, the true IO curve is computed by fitting the sigmoid model (1) to 100,000 stimulus-response pairs. Figure 3-a shows stimulus-response pairs and computed true IO curve for a representative run. In the IO SCE settings, twenty samples are taken from the baseline. The maximum pulse strength is set to 100%. The stopping rule is checked for *T* = 5 as previously recommended [36].

**Fig. 3:**
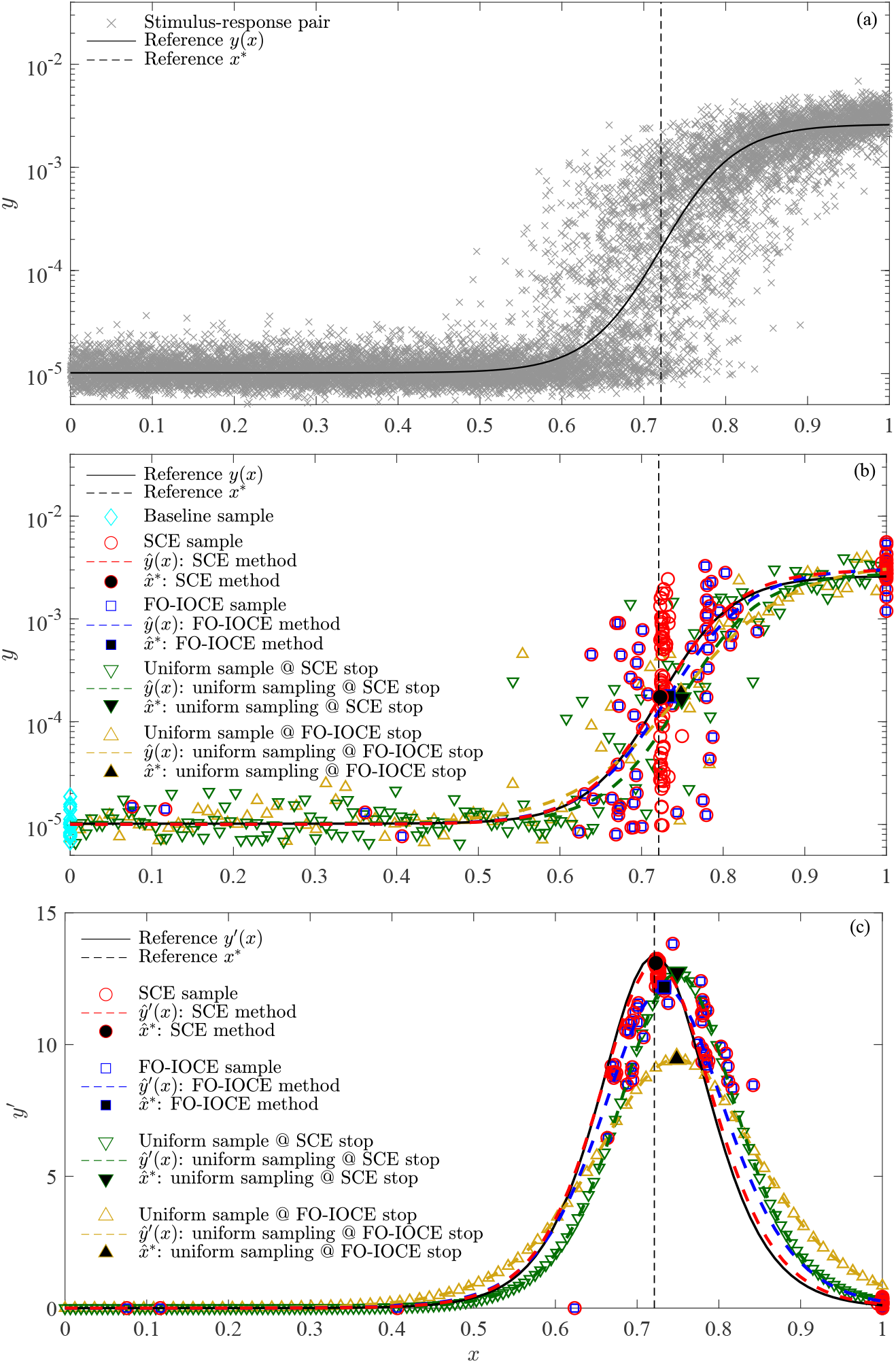
Sample simulation run; a) reference data and IO curve, b) the acquired samples and estimated IO curves by using FO-IOCE method [36] when it stops at 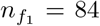, by using the proposed SCE method when it stops at 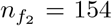, and by uniform sampling method at 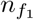 and 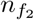, c) The corresponding estimated slope curves. The estimated points of the peak slope (*x*\*, *y ′* (*x*\*)) are shown by filled markers. The x-axis is normalized TMS pulse amplitude, peak-to-peak amplitude of the MEPs is employed in this paper.

The trust-region algorithm serves for the curve fittings, searching for the optimal estimate of the parameter vector ***θ*** between a lower bound 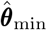 and an upper bound 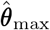 per

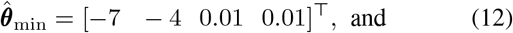

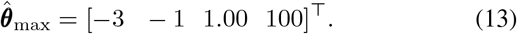

In order to mitigate the variabilities’ effects, the curve fitting is performed on the logarithmic scale [46], [47]. A bad-fit detection and removal technique is employed to improve the performance of the methods.

The maximum number of TMS pulses, *N*_max_ is set to 200 for several reasons. The results in [36] show that the FO-IOCE stopping rule is satisfied with a median of 88 TMS pulses in more than 92% of 10,177 simulation cases. Thus, *N*_max_ = 200 is sufficiently greater than this value, and it is expected that the majority of the runs satisfy the stopping rule before reaching the maximum number of TMS pulses. In practice, there must also be a delay between two pulses, from 5 s to 10 s, thus, applying 200 pulses as well as the required computational process need at least more than 20 minutes, which is long enough from the clinical perspective.

For comparison studies and after stimulating each TMS pulse and completing the IO SCE computational processes, the parameter vector is also estimated by using the same numbers of TMS pulses uniformly distributed between 0 and 100%.

For both the IO SCE and uniform sampling methods, stimulated TMS pulses and their MEP responses, estimations of the parameter vector, as well as the relative errors, all are recorded for comparative studies. The relative estimation errors of the parameters, after applying *n* TMS pulses, *n* = 1, 2, …, are computed and recorded as follows:

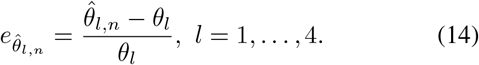

The relative estimation error of IO curve derivative at *x*\*, after applying *n* TMS pulses is computed and recorded as follows:

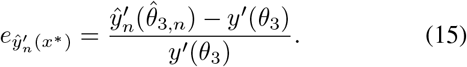

For the IO SCE, 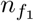 and 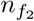 are recorded, denoting when the stoppling rule is satisfied for the first and second times, respectively. For uniform sampling, if the stopping rule is satisfied for *T* = 5, the successful termination of the uniform sampling method is reported and recorded.

### A. A representative run

Figure 3 shows the performance of the proposed IO SCE and uniform sampling methods for a representative run. Figure 3-a shows 100,000 stimulus–response pairs generated by using the statistical model [43]. The reference IO curve is obtained by fitting to this data set. After acquiring the baseline data and the stimulus–response pairs of the initial three pulses, the next pulse intensities are chosen by solving the optimization problem (8) until the stopping rule is satisfied for the first time at 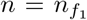. For this representative run, 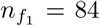. The FO-IOCE method stops at 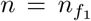, and does not proceed. The proposed IO SCE method continues stimulation by apply TMS stimuli for 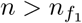 by using (11) until the stopping rule is satisfied for the second time at 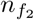. The proposed IO SCE stops at this point. For this representative run, 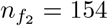.

Figure 3-b shows the acquired samples and estimated IO curves by using FO-IOCE method [36] when it stops at 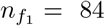, by using the proposed SCE method when it stops at 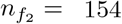, and by uniform sampling method at 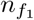 and 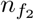.

Figure 3-c shows the corresponding estimated slope curves. The estimated points of the peak slope (*x*\*, *y ′* (*x*\*)) are shown by filled markers. For a better visualization of the IO SCE sampling strategy, the map of stimulus-response pairs are also plotted in the derivative plane.

It is seen that the FO-IOCE method [36] administers stimulus amplitudes, mainly from Regions 2, 3, and 4. The proposed IO SCE method also administers stimulus amplitudes from Regions 5 which significantly improves the slope curve estimation for this run.

It is also seen that the uniform sampling, administers many TMS stimuli from unimportant areas, which do not improve the slope curve estimation.

Figure 4 shows the estimated parameter vector ***θ*** = [*y*^*l*^, *y*^*h*^, *m, s*]^⊤^ for the representative run, for the FO-IOCE method [36] up to its stop at 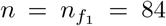, and for the proposed SCE method up to its stop at 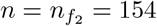, and for the uniform sampling method up to 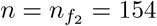. It is seen that the proposed IO SCE method improves the estimation of the mid-point and slope parameters compared to the FO-IOCE and uniform sampling techniques.

**Fig. 4:**
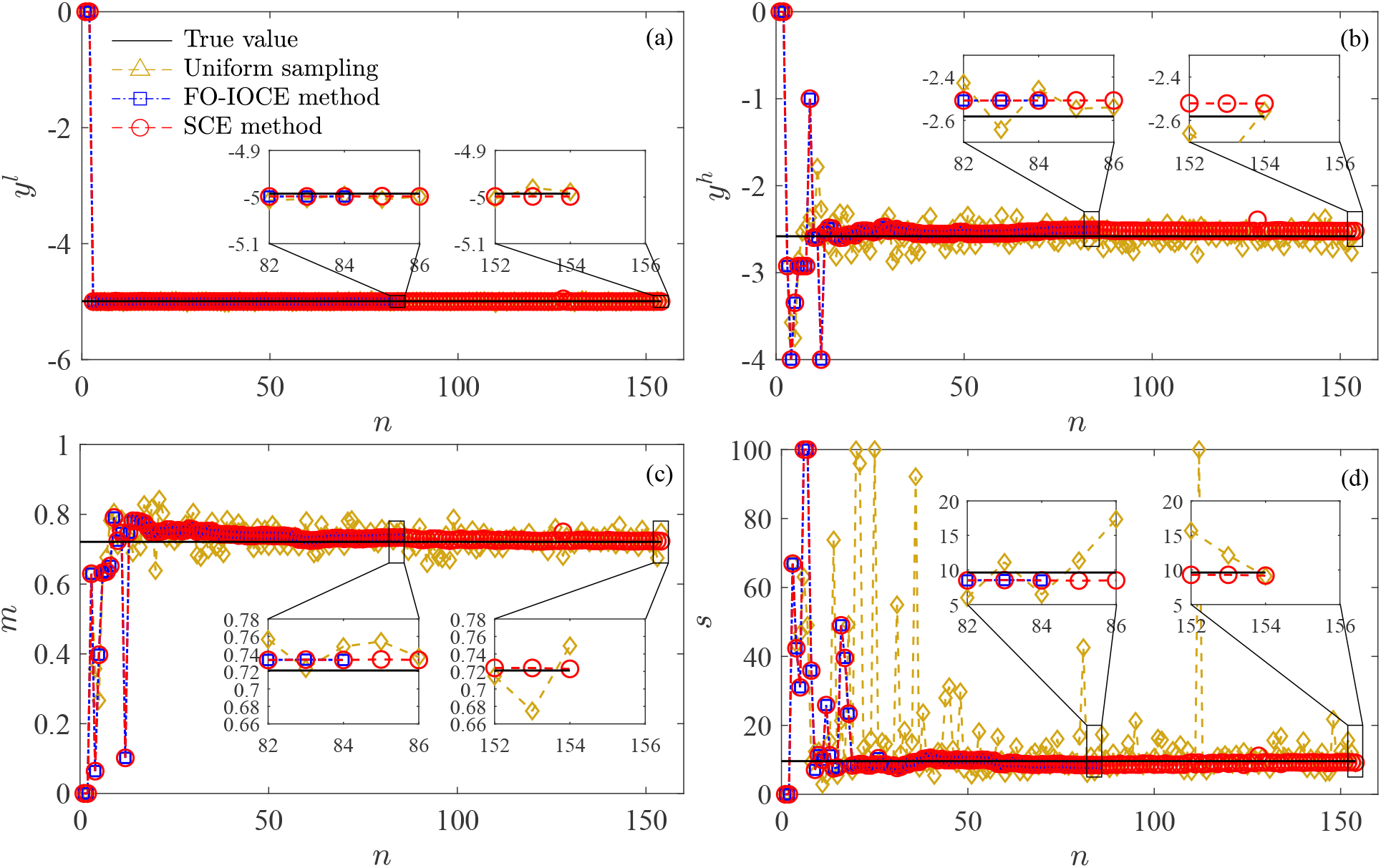
Estimated IO curve parameters for the representative run for the FO-IOCE method stopping at 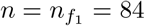, for the proposed IO SEC method stopping at 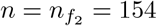, all compared to the uniform sampling method. (a) low plateau *y*^*l*^, (b) high plateau *y*^*h*^, (c) midpoint *m*, and (d) slope parameter *s*.

### B. Comparison between the proposed IO SCE, FO-IOCE, and uniform sampling methods

1. *In terms of the relative estimation errors:* Figure 5 shows the absolute relative estimation errors (AREs) of the parameter vector ***θ*** = [*y*^*l*^ *y*^*h*^ *m s*]^⊤^ for the proposed IO SCE method compared to FO-IOCE and uniform sampling over 1000 runs. The mean, 5th/95th percentiles, and their values are shown in the figure. Except for the low-side plateau *y*^*l*^, IO SCE improves the 95th percentile and mean values as follows:

**Fig. 5:**
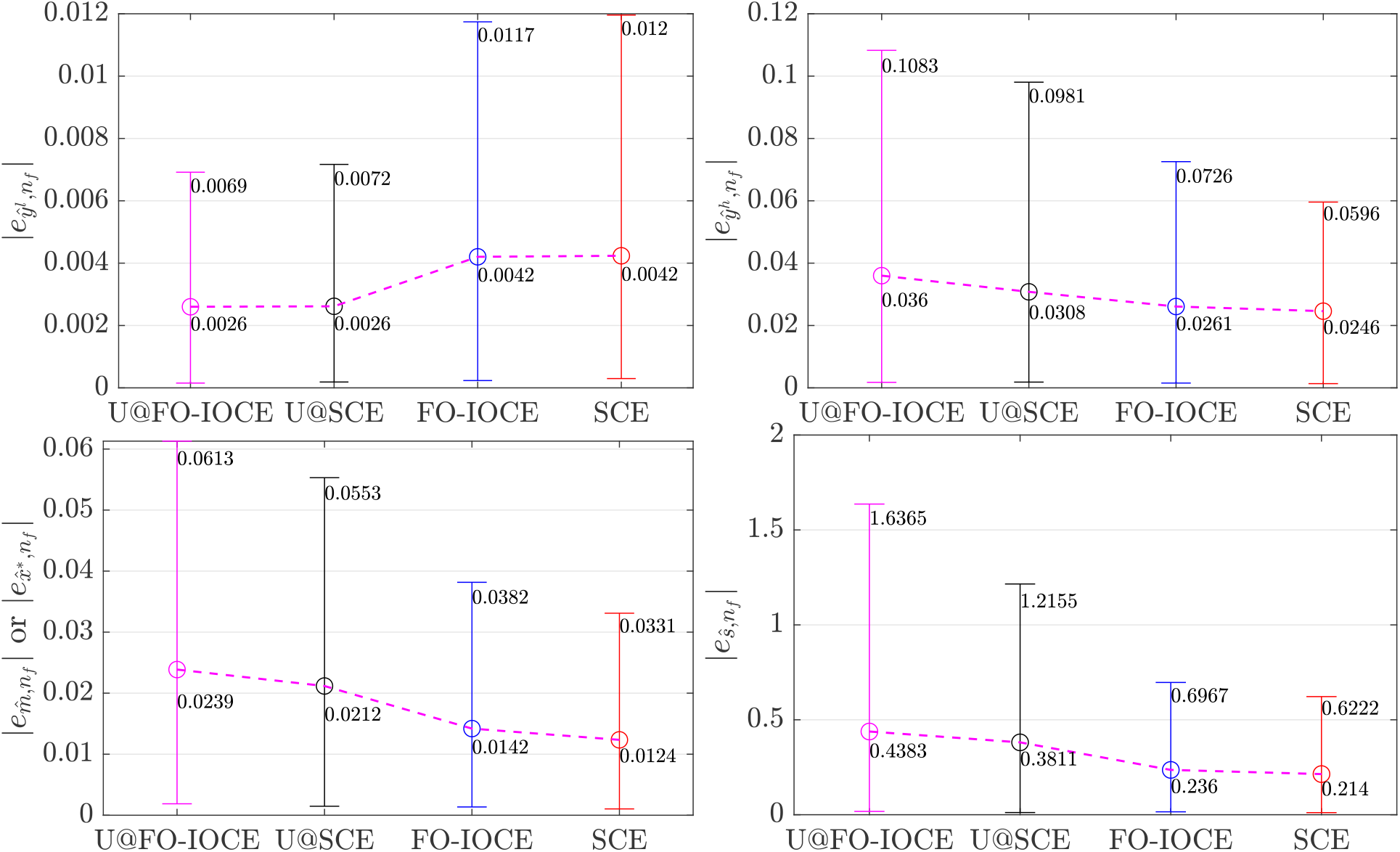
Absolute relative errors for the FO-IOCE and proposed SCE methods at their successful stopping iterations, and compared to the uniform sampling methods. (a) low plateau *y*^*l*^, (b) high plateau *y*^*h*^, (c) midpoint *m*, and (d) slope *s*. Markers and whiskers show the mean and 5th/95th percentile values, respectively. The x-axis tick labels denote the followings. The successful stop occurs when the stopping rule is satisfied before applying the maximum number of TMS pulses, i.e., the *N*_max_-th TMS pulse. The x-labels U@FO-IOCE and U@SCE represent the results of uniform sampling technique at the successful stops of the FO-IOCE and SCE methods, respectively.
  i. The IO SCE method at the time when it successfully stops, results in smaller 95th percentiles and means of AREs compared to the uniform sampling method for *y*^*h*^, *m*, and *s*. When the proposed IO SCE method successfully terminates, the 95th percentiles (mean value in parentheses) of the AREs of *y*^*h*^, *m*, and *s* are reduced by 3.85% (0.62%), 2.22% (0.88%), and 59.33% (16.71%), respectively, compared to those of uniform sampling.
  ii. When the IO SCE method successfully terminates, the 95th percentiles (mean value in the parentheses) of the AREs of *y*^*h*^, *m*, and *s* are reduced by 1.3% (0.15%), 0.51% (0.18%), and 7.45% (2.2%), respectively, compared to those of the FO-IOCE method at its stopping point.
  iii. If the uniform sampling continues until the stopping point of the proposed SCE method, the 95th percentiles (mean value in the parentheses) of the AREs of *y*^*h*^, *m*, and *s* are reduced by 1.02% (0.52%), 0.6% (0.27%), and 42.1% (5.72%), respectively, compared to those of the uniform sampling method which stops at the stopping point of the FO-IOCE method. The AREs of the low-side plateau *y*^*l*^ in all methods are small, with the 95th percentiles below 1.2%. However, uniform sampling shows smaller AREs of *y*^*l*^ than those in the FO-IOCE and SCE methods. This is a result of only 20 samples taken from the baseline in this simulation study. By increasing this value for instance to 100 (as per [36]), the same AREs could be achieved for all these methods. Note that the baseline data mainly effects the estimation of *y*^*l*^. Figure 6 shows the AREs of the IO slope curve at *x*\*. When the proposed IO SCE successfully terminates, its 95th percentile (mean value in the parentheses) of |*e*_*ŷ*′_ (*x*\*)| is reduced by 46.29% (13.13%) compared to that of the uniform sampling method, and by 9.96% (2.01%) compared to that of the FO-IOCE method at its stopping iteration. The 95th percentile (mean value in the parentheses) of |*e*_*ŷ*′_ (*x*\*)| of the uniform sampling method at the stopping iteration of the SCE method is also reduced by 46.25% (6.02%) compared to that of uniform sampling at the stopping iteration of the FO-IOCE method.

**Fig. 6:**
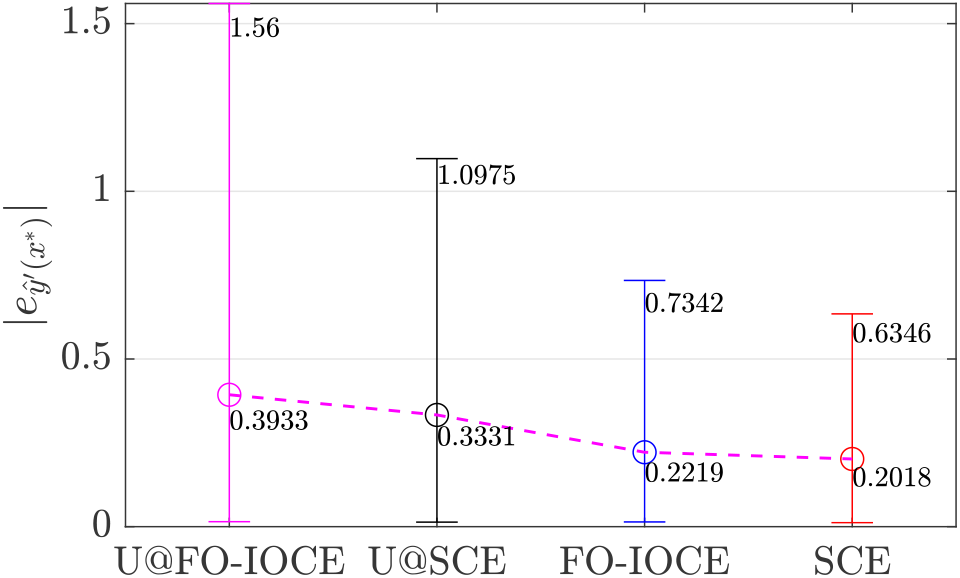
Absolute relative errors of the IO slope curve at *x*\*, for the FO-IOCE and SCE methods at their stopping iterations, compared to the uniform sampling method. The x-labels U@FO-IOCE and U@SCE represent the results of uniform sampling technique at the successful stops of the FO-IOCE and SCE methods, respectively.
2. *In terms of successful termination & number of stimuli:* In 795 (79.5%) out of 1000 runs, the IO SCE method satisfies the stopping rule with an average of 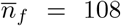 stimuli. Compared to the FO-IOCE method [36] which successfully stops at 866 (86.6%) of 1000 runs with the average of 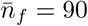 stimuli, the proposed SCE requires 18 more TMS stimuli on average. However, the proposed IO SCE method significantly improves the estimation performance compared to the FO-IOCE method. In none of the 1000 runs, the uniform sampling satisfies the stopping rule prior to *N*_max_ = 200, providing no information about the number of samples for the estimation of the IO slope curve.

Overall, the results confirm that the proposed IO SCE improves the estimation performance at the time when it successfully stops, compared to the FO-IOCE and uniform sampling methods.

## V. Discussions, Conclusions and Future Work

This paper proposes a novel method for input–output slope curve estimation (IO SCE) in transcranial brain and spinal stimulation, such as TMS and TES. The proposed IO SCE is based on sequential parameter estimation (SPE) and optimal sampling principles using Fisher information matrix (FIM) optimization but focusses on the slope curve. The results confirm that Fisher-information-based optimal IO cruve estimation (FO-IOCE) method with subsequent derivation is not optimal for the slope curve estimation, since different inputs (from different regions) are required for the optimal estimation of sigmoidal and normal functions. The results again highlight significant drawbacks of the uniform sampling method in the estimation of the TMS IO slope curve from various perspectives. The uniform sampling’s estimation variation is very large, and none of the runs could satisfy the same stopping rule used for the proposed IO SCE method. Thus, it is not clear how many TMS pulses should be administered for the estimation. In addition, the absolute relative errors of the uniform sampling method are significantly larger than those achieved by the IO SCE method.

As discussed in the paper, there is no tool for the direct measurement of the IO slope at a given pulse strength. Thus, a method was proposed, which firstly estimates the slope curve’s parameter vector through curve fitting in the IO curve domain, and then computes the slope curve using the derivative equation (4). The proposed IO SCE method is sub-optimal in the sense of sampling strategy. Further research is required for the development of a full optimal IO SCE.

The test data presented a wide range of different slopes, x shifts, and intra-individual trial-to-trial variability levels to the studied methods. The realistical model allowed testing with a high number of cases beyond what a reasonable experimental test could and, more importantly, extract ground truth information. The tails of the incorporated probability density functions and extreme-value distributions of the model furthermore challenged the methods also with rare extreme cases, which may not be well represented in typical study cohort sizes. The IO SCE method is ready to perform with most available devices and primary motor cortex representations of many muscles as they mostly affect the above three parameters of slope, x shift, and trial-to-trial variability [48]. The use of differently strong pulse sources, for instance, primarily scales the x axis, resulting in a scaled slope [47], [49], [50]. The focality of the coil might affect recruitment and therefore slope as well as trial-to-trial variability, but there is little quantitative information on that aspect in TMS yet [51]–[53]; both clinical procedures and safety guidelines assume that focal and unfocal coils are rather comparable with respect to MEPs and motor threshold [54]. Suboptimal placement of stimulation coils, on the other hand, leads to a lower electric field at the target and thus scaling of the x axis similar to weaker pulse sources [55], [56]. In addition, it may change the trial-to-trial variability due to (modulatory) interaction of the brain areas co-activated with the intended target. Latest experiments with deep-brain electrical stimulation of the corticospinal tract suggests similar statistical properties of slope, x shift, and particularly the individual contributions to the trial-to-trial variability so that the method might readily translate to such applications as well [57].

The presented work has focused on the MEP’s peak-to-peak amplitude, while the area under the curve as an amplitude estimate could be used likewise. Furthermore, this work intends to stimulate and inform measurement procedures for other MEP-related qualities, such as response latency or cortical silent period, for a faster and more accurate estimation. Clinical tests, and neurophysiological studies by using the proposed IO SCE method are suggested for future research.

## Appendix A

### Matrices IN FIM Optimization Problem per Eq. (8)

The matrix 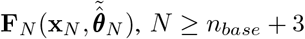, is computed as

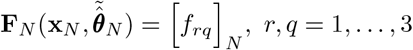

where

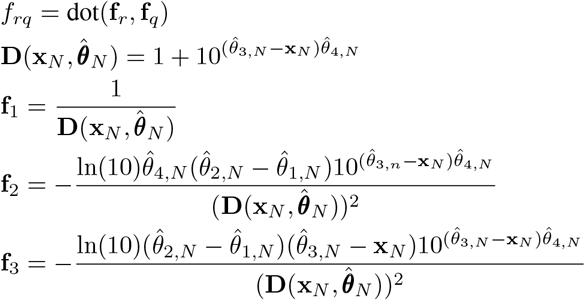

dot(**f**_*r*_, **f**_*q*_) returns the scalar product of the vectors **f**_*r*_ and **f**_*q*_.

The matrix 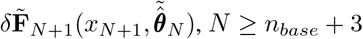 is computed per

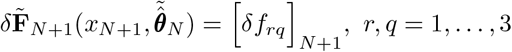

where

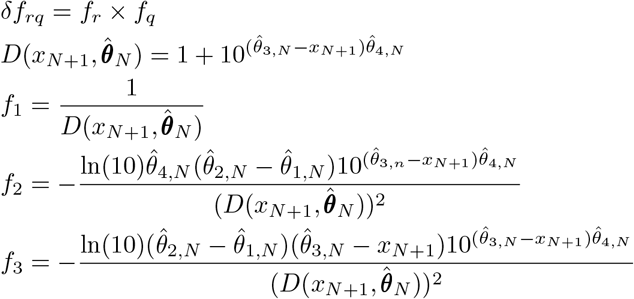

## Abbreviations & Nomenclatures

In this paper, vectors and matrices are written in bold letters. ***θ***^⊤^ denotes the transpose of the vector ***θ***. The operator ‘^’ denotes estimations, e.g., 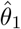 refers to the estimation of *θ*_1_. The overline signifier ‘—’ is used for the average values, e.g., 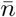 denotes the average of *n*.

TM: Transcranial magnetic stimulation
IO: Input–output
MEP: Motor evoked potential
FIM: Fisher information matrix
FO-IOCE: Fisher-information-based optimal IO curve estimation
SCE: Slope curve estimation
ARE: Absolute relative estimation error
*x*: TMS pulse strength normalized between 0 and 1, *x* = 1 means 100% pulse strength
*x*_min_: Minimum TMS pulse strength. It is presumed to be 0 in this work
*x*_max_: Maximum TMS pulse strength. It is presumed to be 100% in this work
*y*(*x*): (Reference) IO curve
*y′* (*x*): (Reference) IO slope curve, *y′* = *dy/dx*
*y*^*l*^: (Reference) low-side plateau of IO curve
*y*^*h*^: (Reference) high-side plateau of IO curve
*m*: (Reference) midpoint of IO curve
*x*\*: (Reference) TMS pulse strength which targets the midpoint of the IO curve, i.e., *x*\* = *m*
*s*: (Reference) slope parameter
*θ*: (Reference) parameter vector ***θ*** = [*θ*_1_ *θ*_2_ *θ*_3_ *θ*_4_]^⊤^ = [*y*^*l*^ *y*^*h*^ *m s*]^⊤^
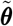: Parameter vector used for FIM optimization 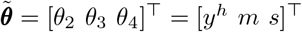
*n*_*base*_: Number of baseline samples
*n*: Number of TMS pulses
*N*: Total number of samples *N* = *n*_*base*_ + *n*
*N*_0_: Size of the data set for the initial curve fitting. It is equal to the number of baseline samples plus the first three initial TMS pulses, *N*_0_ = *n*_*base*_ + 3
*N*_max_: Maximum number of TMS pulses
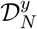: IO data set used for curve fitting
**x**_*N*_: All TMS stimuli, 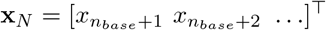
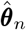: Estimation of ***θ*** after the stimulation of the *n*-th TMS pulse
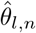: Estimation of *θ*_*l*_, *l* = 1, 2, 3, 4, after the stimulation of the *n −* th TMS pulse
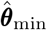: Lower bound of the estimate of ***θ***, i.e., estimated parameters cannot be lower than values in 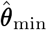
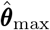: Upper bound of the estimate of ***θ***, i.e., estimated parameters cannot be greater than values in 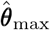
*ŷ*_*n*_: Estimation of the IO curve after the stimulation of the *n −* th TMS pulse
*ϵ*_*l*_: Convergence tolerance
*T*: Number of successive times the convergence criterion must be satisfied in stopping rule
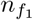: Number of TMS pulses satisfying the stopping rule for the first time
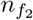: Number of TMS pulses satisfying the stopping rule for the second time. The proposed IO SCE method successfully stops at this point
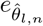: Relative estimation error of *θ*_*l*_, *l* = 1, 2, 3, 4, after the stimulation of the *n−*th TMS pulse
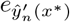: Relative estimation error of IO curve derivative at *x*\*, after the stimulation of the *n −*th TMS pulse

